# Sequencing of SARS CoV2 in local transmission cases through oxford nanopore MinION platform from Karachi Pakistan

**DOI:** 10.1101/2021.01.07.425705

**Authors:** Samina Naz Mukry, Shariq Ahmed, Ali Raza, Aneeta Shahni, Gul Sufaida, Arshi Naz, Tahir Sultan Shamsi

## Abstract

The first case of severe acute respiratory syndrome 2 (SARS CoV2) was imported to Pakistan in February 2020 since then 10,258 deaths have been witnessed. The virus has been mutating and local transmission cases from different countries vary due to host dependent viral adaptation. Many distinct clusters of variant SARS CoV2 have been defined globally. In this study, the epidemiology of SARS CoV2 was studied and locally transmitted SARS CoV2 isolates from Karachi were sequenced to compared and identify any possible variants.The real time PCR was performed on nasopharyngeal specimen to confirm SARSCoV2 with Orf 1ab and E gene as targets. The viral sequencing was performed through oxford nanopore technology MinION platform. Isolates from first and second wave of COVID-19 outbreak in Karachi were compared. The overall positivity rate for PCR was 26.24% with highest number of positive cases in June. Approximately, 37.45% PCR positive subjects aged between 19-40 years. All the isolates belonged to GH clade and shared missense mutation D614G in spike protein linked to increased transmission rate worldwide. Another spike protein mutation A222V coexisted with D614G in the virus from second wave of COVID-19. Based on the present findings it is suggested that the locally transmitted virus from Karachi vary from those reported from other parts of Pakistan. Slight variability was also observed between viruses from first and second wave. Variability in any potential vaccine target may result in failed trials therefore information on any local viral variants is always useful for effective vaccine design and/or selection.

**Author’s summary:** Despite precautionary measures the COVID-19 pandemic is causing deaths all over the world. The continuous mutations in viral genome is making it difficult to design vaccines. Variability in genome is host dependent and data sharing has revealed that variant for different geographical locations may harbor different mutations. Keeping this in mind the current study was focused on the epidemiology of SARS CoV2 in symptomatic and asymptomatic COVID –19 suspected cases with impact of age and gender. The locally transmitted SARS CoV2 isolates from Karachi were sequenced to compared and identify any possible variants. The sequenced viral genome varied from the already submitted sequences from Pakistan thereby confirming that slightly different viruses were causing infections during different time periods in Karachi. All belonged to GH clade with D614G, P323L and Q57H mutations. The virus from second wave had A222V mutation making it more different. This information can be useful in selecting or designing a vaccine.

## Introduction

Corona virus disease (COVID-19) is a transmissible infectious disease caused by a newly emerged beta corona virus - SARS CoV2 originated as a result of viral spill over from animals[1]. It is a positive sense, enveloped, single stranded RNA virus of genus betacoronavirus [2]. The exact origin of this virus either from bat, pangolin or any other mammal is still under debate[1,3]. Metagenomic analysis of SARS Cov2 has revealed that it a distinct virus very closely related to SARS CoV. COVID-19 pandemic started in December 2019 after first reports from Wuhan China[3,4]. To date it has affected 188 countries with a death rate of 2.31%. The death rate for SARS CoV2 infection is lower than SARS CoV with a higher transmission capability [5]. The global elderly population was most affected with higher mortalities due to acute respiratory distress syndrome(ARDS). The first SARS Cov2 genome was sequenced and published in December 2019 by Wang *et al*[6]. The genome of SARS CoV2 is ~29.9 kb long with orf1ab at the 5’end and spike protein(S), envelope protein (E) and matrix protein (M) coded at the 3’ end. Seven viral accessory proteins are also coded by *ORF3a, ORF6, ORF7a, ORF7b, ORF8a, ORF8b* and *ORF10* genes[7]. The virus has sixteen non-structural proteins (NS1-NS16). The infection initiates with lower respiratory discomfort which progress to pneumonia often causing sudden deaths [4,8]. The virus establishes itself by binding through receptor binding spike protein to angiotensin-converting enzyme 2 (ACE2) receptors in lungs [4]. Data from various studies support that a virus induced excessive exaggerated immune reaction or cytokine storming extensively damage tissues ACE2 receptors expressing organs in the host [2].

In the recent outbreak of COVID-19 asymptomatic carrier were capable of transmitting virus to healthy human. As reported earlier, during SARS CoV outbreak in 2002/2003 variant viruses evolved due to possible transformation events within host [9]. Likewise, since its first report many mutations have been reported in SARS CoV2. A large number of SARS CoV2 sequences have been deposited in respective repositories since the beginning of the pandemic[10,11]. Due to relatively lower mutation rates different branches or clades have been defined[12]. The clinical significance of all these clades is yet to be defined. As per Nextstrain an open source project so far five large clades of SARS CoV2 (19A, 19B, 20A, 20B and 20C) have been identified all over the world. The earlier strains identified in 2019 belonged to clade 19A and 19B. The clade 19B (GISAID S) differed from the root clade 19A by substitutions of C>T and T>C at positions 8782 and 28144 respectively. The clade 20 A had unique substitutions of C>T at positions 3037 and 14408 along with substitution G>C28883. The clade 20B originated with consecutive substations 28881G>A, 28882G>A, and 28883G>C whereas 20C has substitutions 1059C>T and 25563G>T. The clades 19A and 19B were geographically linked to Asia. The clades 20A and 20B were prevalent in Europe therefore seemed to originate from there. The prevalent strain in North America was the clade 20C in later half of 2020. The SARS CoV2 sequences submitted from Pakistan belonged to clades 19A, 19B, 20A and 20B. Rapid whole genome sequencing coupled with prompt data sharing is the key to understand the emergence of variants and geographic epidemiology of SARS CoV2 variants during current pandemic[13]. Clinical correlation of these variants with disease transmission dynamic, treatment responsiveness and fatality rates have also been extensively studied[14]. Among the various sequencing platforms third generation sequencing of SARS CoV whole genome by Oxford nanopore MinIon technology based sequencing has gained popularity. The advantage of this plat form is that long reads of virus genome are obtained and time to data acquisition and analysis is also reduced as compared to other methodologies [15]. The objective of present study was to evaluate the epidemiology of SARS CoV2 in symptomatic and asymptomatic COVID –19 suspected cases with impact of age and gender. Furthermore, locally transmitted SARS CoV2 isolates from Karachi were sequenced to compared and identify any possible variants.

## Results

Two thousand and sixty five (2065) PCR tests were performed from May to November 2020 at COVID-19 Lab. of NIBD. The overall positivity rate for PCR was 26.24%. Highest number of positive cases with increased viral load (lower Ct values) was observed during the month of June (Fig 1a). The Ct value for SARS CoV2 ranged between 10.8 to 34.32 in June with a median Ct value of 24.2 (Fig 1b). After the first wave of COVID-19 in Karachi an increase in positive cases was observed in October; after a decline in August and September; with the lowest median Ct value (20.21) between May to November (Fig 1a). A large number of patients were negative for SARS CoV2 with COVID-19 like symptoms caused by probably some other viral or bacterial infection. The commonest symptom was weakness for PCR positive and PCR negative groups. The frequency of SARS CoV2 positive males (27.5%) was slightly higher than females (26.28%). Approximately 37.45% PCR positive subjects aged between 19-40 years.

**Fig 1:**
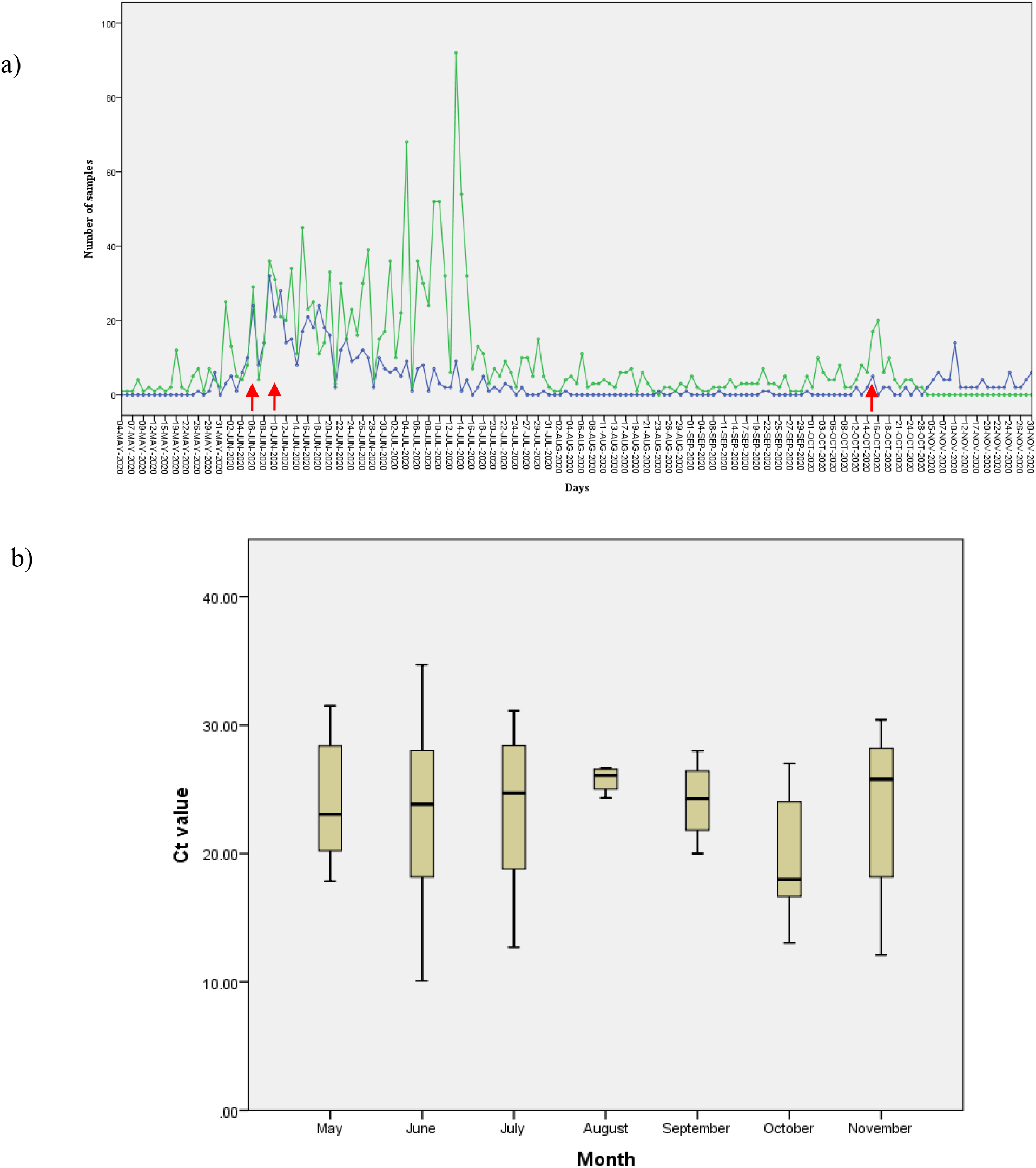
Detailed overview of SARS CoV2 suspected samples analyzed at National Institute of Blood Diseases and Bone Marrow Transplantation. Nasopharyngeal swabs were tested for presence of SARS CoV2 RNA using commercial kit. X axis (horizontal) shows dates from 4^th^ May to 30^th^ November 2020 when samples were taken. A) Graphical summary of tested specimen per day. The green line shows samples negative SARS CoV2 per day and the blue line shows positive samples. The dots indicate total samples for each day. The red arrow indicates the date of collection for samples 1= NIBD01-PAK-KHI, 2=NIBD02-PAK-KHI, 3=NIBD03-PAK-KHI, NIBD04 selected for SARS CoV2 whole genome sequencing. B) The box and whiskers graph where Ct median value with SARS CoV2 specific Orf1ab as target is indicated by line and the box is extended from 25th to the 75th percentiles. Lowest median Ct value 20.21 was observed in October.

### Viral whole genome sequencing

Four samples with low Ct values <20 (range: 10.08-19.69) were selected. Of the four sequenced samples two were isolated from asymptomatic while other two from mild or moderate COVID-19 patients each. The age range of selected cases was 22-58 years. Full viral genome sequences from these confirmed cases of local transmission of SARS CoV2 from Karachi during June (peak of first wave) and October (initial phase of second wave) were obtained through MinIon ONT platform. The sequencing details with GISAID accession numbers are listed in Table 1. The genome size obtained was 29903 with depth of coverage between 2976-3653 (Table 1).

**Table 1:** Detailed genomic features of four SARS CoV2 viruses

| Sample | Genbank<br>Accession<br>Number | GISAID<br>Accession<br>No. | Lineage | No of<br>reads | genome<br>Size | GC<br>content | coverage X | Depth of<br>Coverage | Mapped<br>reads |
| --- | --- | --- | --- | --- | --- | --- | --- | --- | --- |
| NIBD01-PAK-KHI | MW4<br>03500 | EPI_ISL<br>_708839 | B.1.<br>36.6 | 19313<br>1 | 29603 | 39% | 100 | 2976 | 98.57% |
| NIBD02-PAK-KHI | MW4<br>00961 | EPI_ISL<br>_709540 | B.1.<br>36.6 | 24023<br>8 | 29836 | 39% | 100 | 2976 | 98.58% |
| NIBD03-Pak-Khi | MW4<br>11960 | EPI_ISL<br>_709544 | B.<br>1.<br>160. | 23288<br>8 | 29594 | 40% | 100 | 3653 | 98.57% |
| NIBD04-Pak-Khi | MW4<br>11961 | EPI_ISL<br>_709542 | B.<br>1.36<br>. | 19790<br>0 | 29519 | 40% | 100 | 3161 | 86.77% |

### Phylogenetic profiling

The whole genome sequences obtained were aligned with the reference genome and 564 global sequences. strains NIBD 01-PAK-KHI and NIBD 02-PAK-KHI clustered with variants from Bangladesh and India with descendent predominantly from Saudi Arabia, India and England. New nodes were defined for NIBD 03-PAK-KHI and NIBD 04-PAK-KHI with divergence of 14 and 27 respectively. Both these were clustered with separate sequences from New Zealand having presumed ancestral connection with isolates from Germany.

The phylogenetic relatedness with previously submitted sequences from Pakistan, India, Saudi Arabia, Netherland, England, New Zealand was observe with a pair wise mean genetic distance of 0.011.

### Mutational analysis

The mutational analysis revealed presence of 21 synonymous mutations, 15 non-synonymous mutations and 2 non-frame shift substitutions altogether, spanning in 5’UTR, spike, orf1ab, orf1a, orf3a, orf7a, orf 8, orf10, N and M protein genes (Table 2). All the four sequences were grouped in clade 20A (GISAID: GH) since characteristic mutation in 5’UTR 241 C>T and nonsynonymous SNV i.e. 3037C>T, 23403A>G (S-D614G), 25563G>T (Q57H) were observed. As per GISAID database four previously submitted sequences from Pakistan also belonged to GH clade but sub-lineage signatures vary with 32 additional mutations observed in the sequences obtained during present study from local transmission cases of Karachi (Table 2). Hence, all the genome sequences were unrelated to the previously reported cluster of SARS CoV2 from Pakistan except for four isolates; 3 from Islamabad and 1 from Kohat belonging to GH clade lineage B (Table 1; Fig 3). The NIBD4-PAK-KHI obtained from a health care worker varied from the other isolates with highest number of mutations (Table 2). An additional clade GV specific nonsynonymous variant 22227C>T (A222V) in spike protein was also present inNIBD4-PAK-KHI along with. About 7% of all GISAID sequences belonged to GV clade which is characterized by presence of this SNV. The virus NIBD4-PAK-KHI is the first variant of GH clade harboring 22227C>T isolated in October from Asia. Close but divergent sequence homology was detected with a variant virus from New Zealand collected in November 2020 (GISAID accession no: EPI_ISL_682284; Fig 3).

**Table 2:**
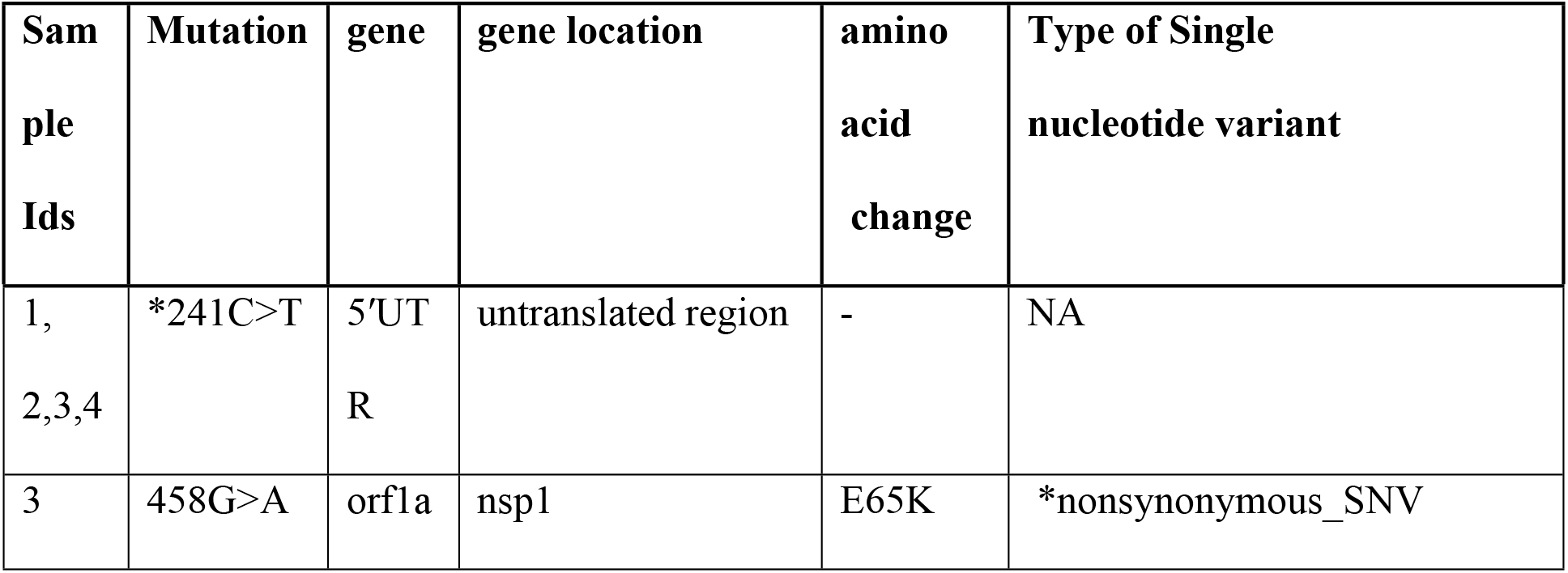
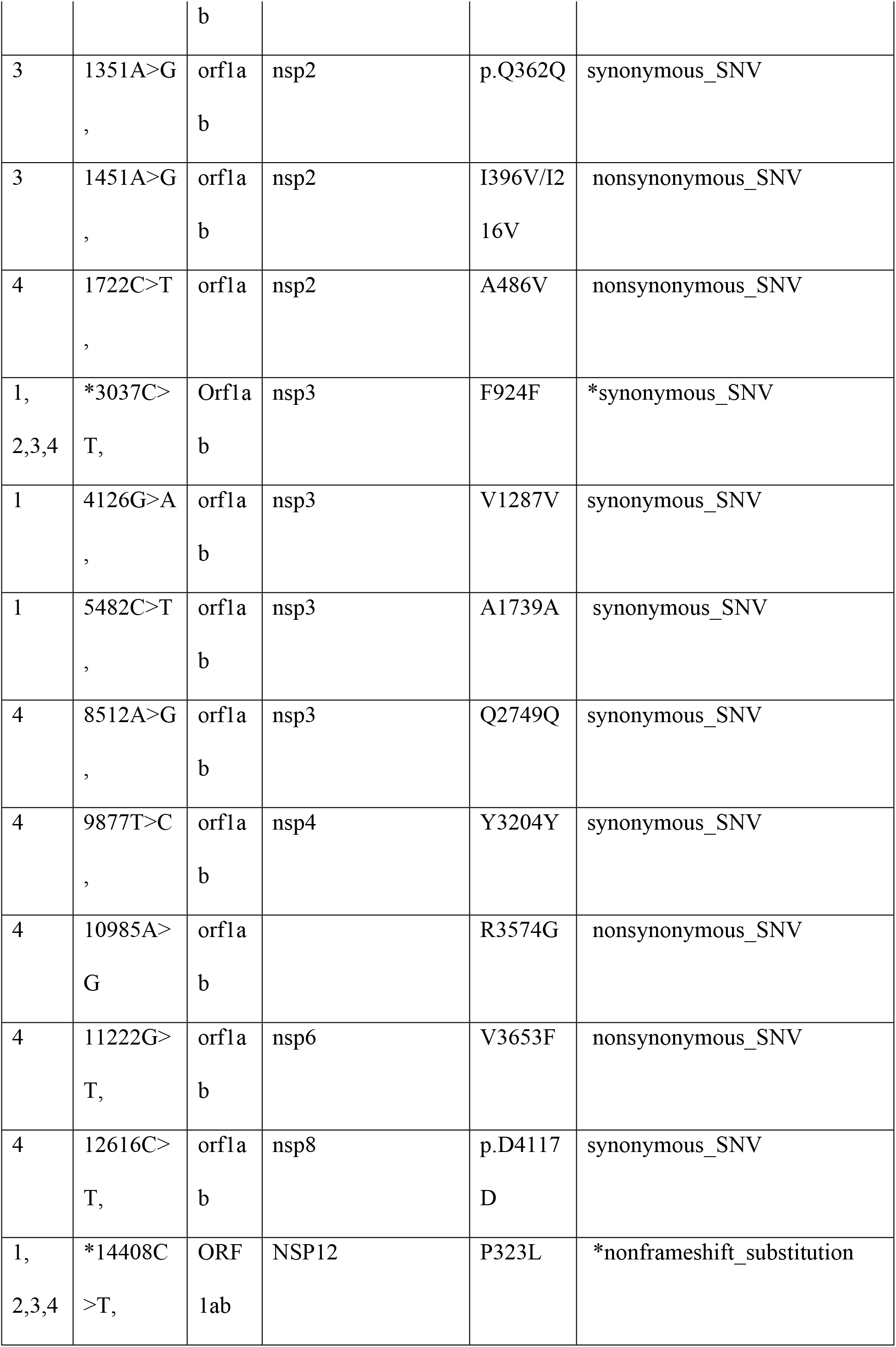
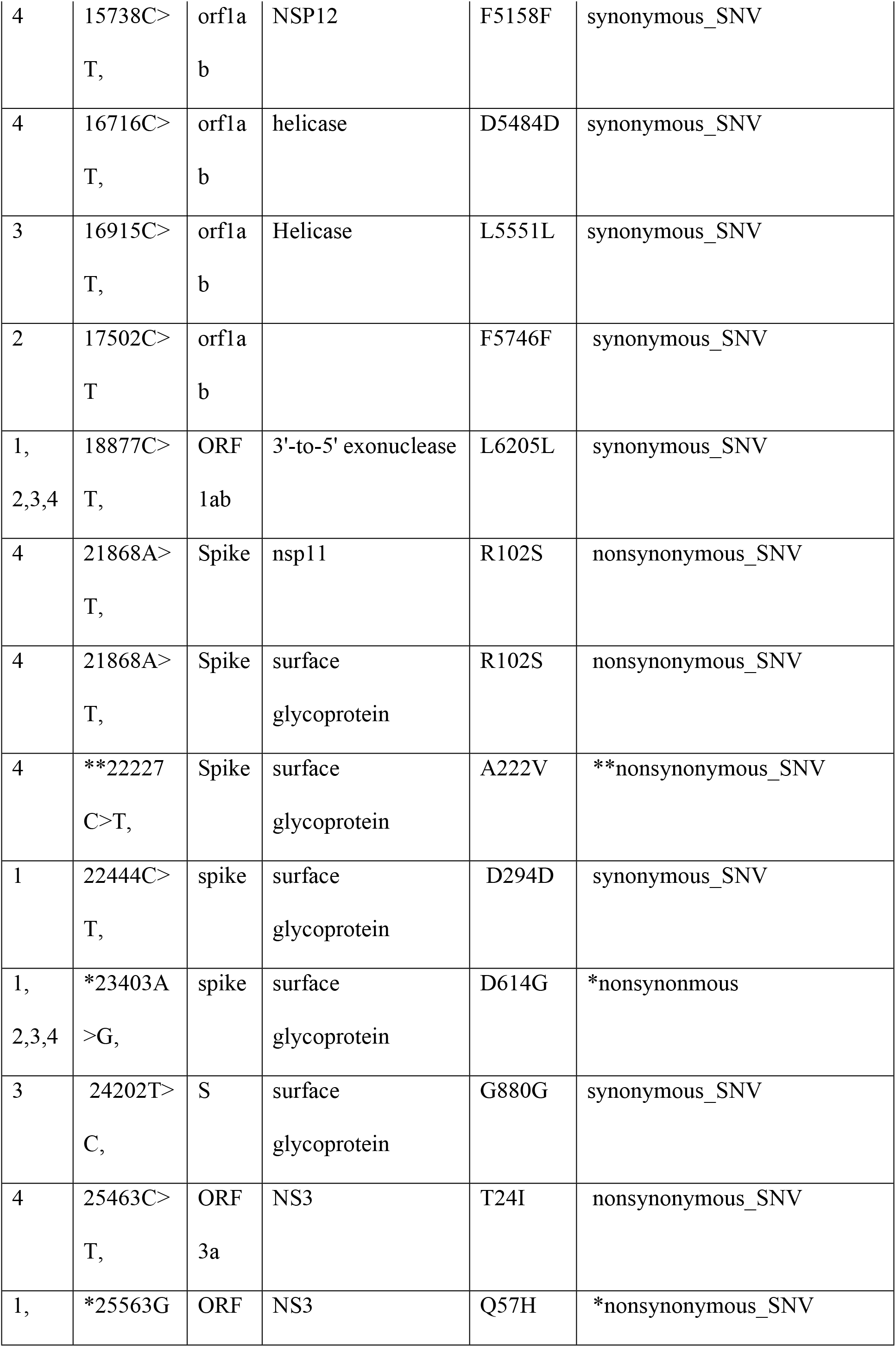
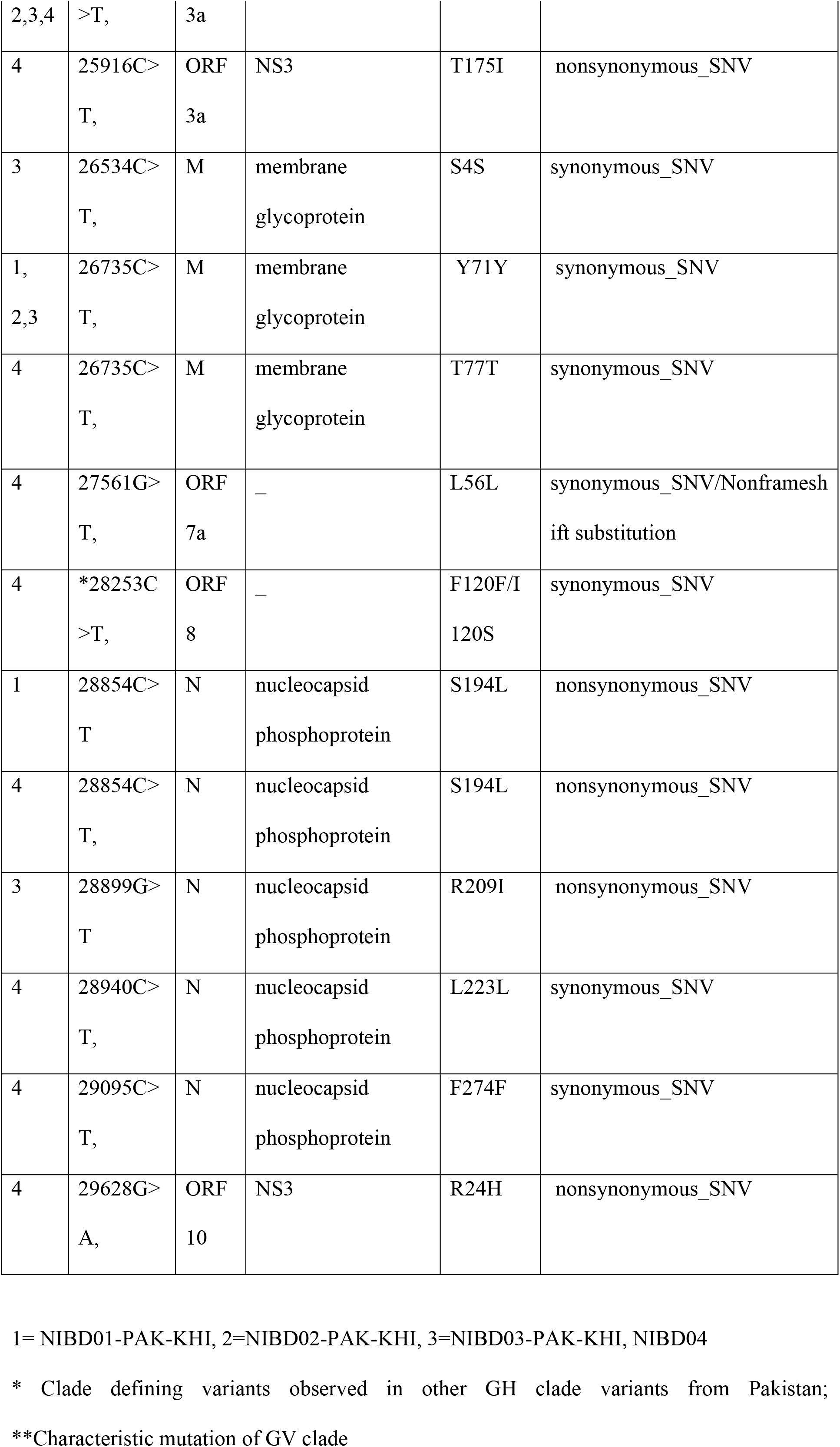
List of single nucleotide variants in genome of locally transmitted SARS CoV2 from Karachi

## Discussion

Karachi is the most populous major metropolitan city of Pakistan. The first imported case of COVID-19 in a returning traveler from Iran was detected in Karachi and the city became the most badly affected area of Pakistan during the first wave of COVID-19. This study is one of the initial report on epidemiological and molecular landscape of SARS CoV2 from Karachi. Based on the retrospective data from NIBD the PCR positivity rate for SARS CoV2 was 24.2% in June (the peak of first wave) which dropped to 5% during September. The positive cases per day increased in October (Fig 1) with a lowest median Ct value observed in October as compared to other months. The increasing trend in PCR positivity correlated well with the regional increase in COVID-19 cases in Sindh. In order to contain SARS CoV2 transmission and also to reduce the possible burden on the striving national healthcare system, the government of Pakistan has officially imposed smart lockdown declaring second wave of COVID-19 with increased cases on 28th October 2020. In order to evaluate any possible genetic variability in community transmitted SARS CoV2 in Karachi four confirmed indigenous SARS CoV2 viruses were randomly selected from the peak of first wave and initial phase of second wave of COVID-19. The genome sequencing was performed through ONT MinIon platform. ONT is claimed to be the most accurate, sensitive and quick method with reduced PCR based bias for detecting minute changes in viral genome. Samples are barcoded to facilitate multiplexing and the tiling generates precise long stretches of genome which could be read easily through defined pipelines thereby decreasing the time for post-acquisition analysis of SARS CoV2 genome [15]. To the best of our knowledge this is the first study from Pakistan on SARS CoV2 genome exploiting the potential of this third generation sequencing method.

The evolution of SARS CoV2 is nonrandom and human host dependent variants are evolving. with over 272,829 virus sequence depositions in GISAID Initiative (epicov.org) alone [16]. SARS CoV2 proteins are heterogeneous with a large number of variable amino acid substitutions with either no or significant impact on viral transmission and transcription within human host [17]. Many prevalent mutations have been defined for SARS CoV2 with signature hotspot mutation for each distinct clade [18]. The lineages for studied viruses were defined through PANGOLINE pipeline and all were placed in lineage B1. with sub lineage differences listed in Table 1. The 23403A>G is a widely documented hotspot mutation in spike protein which replaces aspartic acid with glycine at position 614 altering the viral antigenic properties [19]. Variants with D416G evade initial immune recognition by host resulting in production of autoantibodies and facilitate higher-transmission, infectivity and case fatality rate (CFR). All the sequenced viruses of current study had this mutation along with Q57H in orf3a gene grouping them in Nextstrain clade 20A (GISAID: GH; Table 2). The coexisting Q57H has previously been reported to reduce the virulence of D614G therefore the prevalence of the GH clade in Pakistan may be a probable reason for low mortality rate (2.1%) for COVID-19 cases during July to September. Furthermore, the Q57H amino acid substitution causes truncation of orf3b gene via introduction of a stop codon at amino acid 13 giving rise to full length orf 3b deficient variants [20,21]. These variants were prevalent in Asian and North American countries, including Saudi Arabia, Indonesia, South Korea, Israel, Egypt, USA and Colombia [5,22,23]. All the Pakistani viruses sequenced between June to November 2020 belonged to GH clade with Q57H substitution (Fig 3). The orf3b is a potential serological target for most vaccines, hence, this serological difference should be taken into account while selecting a vaccine for SARS CoV2 in Pakistan. Another prevalent mutation in Pakistani GH isolates is non-frameshift substitution P323L in NSP12 which was linked to higher severity [20]. Almost a 100% coexistence of D614G, P323L and C241T has been reported which is in line with the present observations. This coexistence positively favors viral replication, infectivity, transmission and manipulation of host machinery[24]. Of the 39 mutations observed in the SARS CoV2 genomes sequenced during the present study; 33 SNVs were not observed in previously submitted sequences from Pakistan defining a separate local cluster for SARS CoV2 virus accumulating within Karachi (Fig 2). It may be because of the host driven genetic drift within the viral genome of locally transmitted SARS CoV2 virus [25].

**Fig 2.**
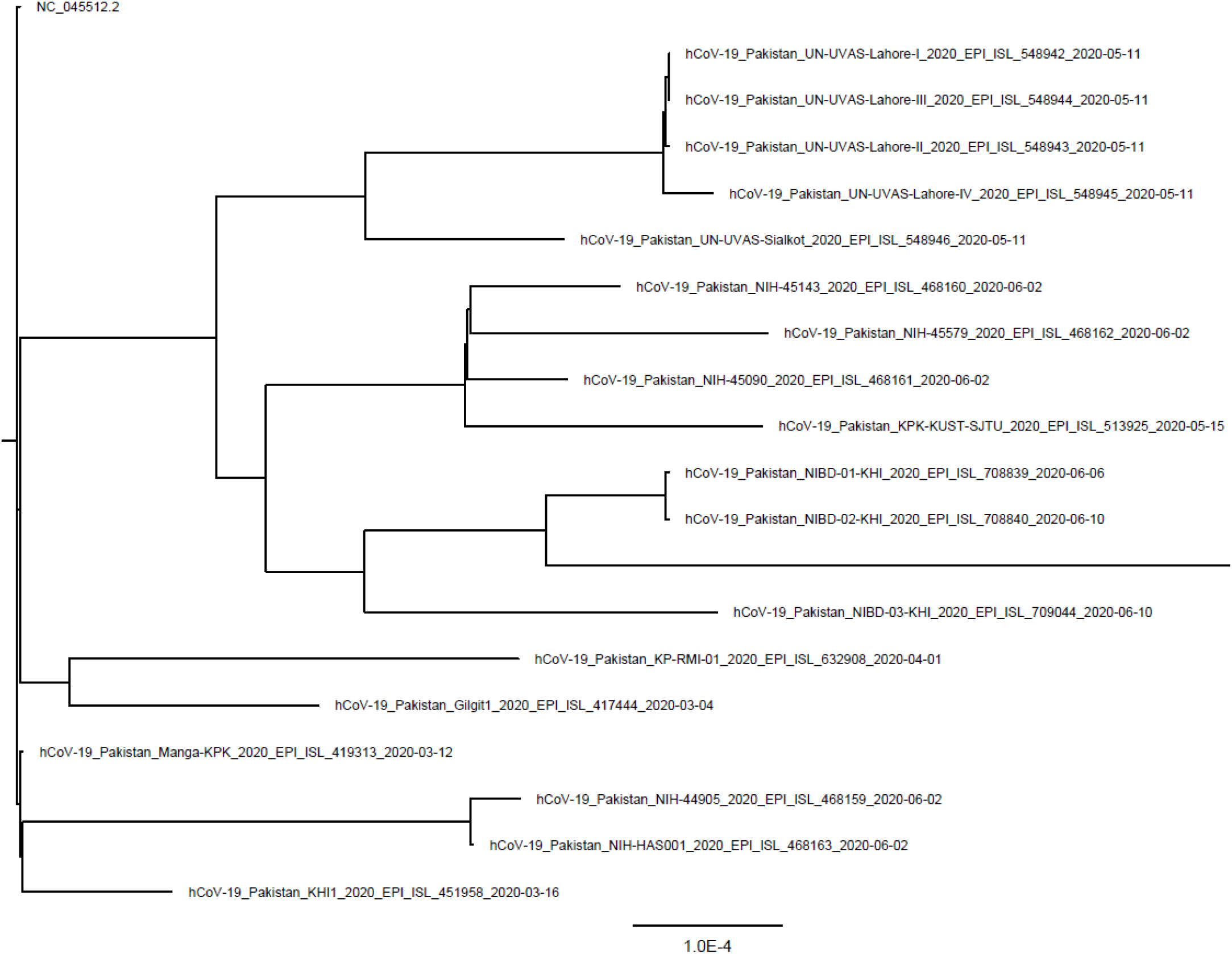
Phylogenetic tree for locally isolated SARS CoV 2 viruses from Pakistan. The locally transmitted SARS CoV2 from Karachi clustered distinctly with viruses from Kohat and Islamabad. The root for the tree is reference sequence from China NC_045512.2

**Fig 3.**
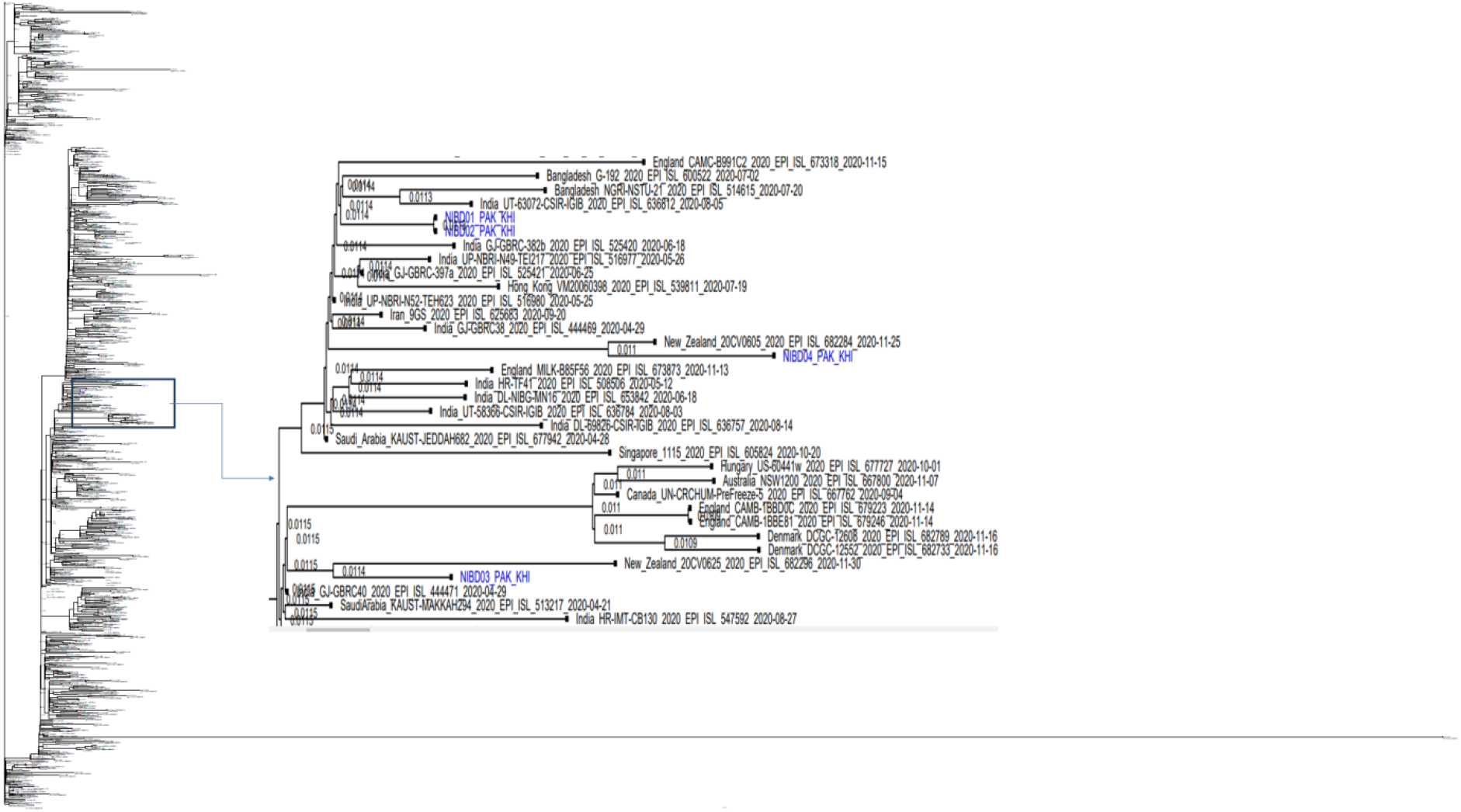
Geo-epidemiological profile of SARS CoV2 viruses from Karachi. The NIBD01-PAK-KHI, NIBD02-PAK-KHI are clustered with viruses from India and Bangladesh. While NIBD01-PAK-KHI and NIBD02-PAK-KHI are clustered with two distinct variants from New Zealand retrieved from GISAID epicov™ The out group for the tree is Wuhan reference sequence NC_045512.2.

Spike A222V mutation was reported from Spain during spring 2020 and clade GV (20A EU1) of SARS CoV2 was defined with increased infectivity and fatality potential [26]. Many mutations overlap between G clade. The global coexistence of A222V mutation within GH clade viruses is rare (0.0015%; GISAID Initiative (epicov.org) with total 84 viruses. The NIBD04 PAK-KHI carried this mutation and is the first sequence from Asia with this unique coexistence sharing ancestral origin from Europe. Within Europe and USA; the A222V is associated with newer waves of COVID-19 with increase viral loads, similarly the virus from the initial phase of second wave of COVID-19 in Karachi was presented with it. It can be assumed that the median viral load corresponding with lower median Ct value may be an indirect indication of accumulation of this virus in Karachi which needs further confirmation via sequencing of virus from the second wave. Another possible reason for this variance in NIBD04 PAK-KHI could be coinfection with two variants of SARS CoV2. The coinfection has been reported with a low prevalence rate of 1.5% in healthcare workers globally. In either case caution is warranted since infectivity potential for this variant is higher throughout Europe.

## Conclusion

Hence it can be concluded that the second wave of COVID-19 may not be clinically distinct but host driven genetic evolution of virus may impact its infectivity and CFR. Future comparative genomic studies with substantial number of virus from first and second waves are suggested in order to understand the possible evolutionary origin, genomic variability and trajectory of anticipated futures waves.

## Methods

This study was conducted at National Institute of Blood Diseases & Bone Marrow Transplantation after approval from NIBD Bioethics committee. A total of 2065 patients of either gender were enrolled. A detailed proforma was filled inorder to obtain the background exposure and brief clinical history of all patients. Sample were collected from nasopharynx of suspected COVID-19 patients and transported in VTM to the laboratory for virus RNA extraction using Favorgene viral RNA extraction kit (Cat# FAVNK 001-2, Favrgene Bio Corp,Taiwan). The real time PCR was performed using CE IVD marked Bosphore Novel Corona virus detection kit (Cat # ABCOW6, Anatolia Geneworks, Turkey) with orf1ab and E genes as viral targets.

### Viral whole genome sequencing

RNA was extracted from fournasopharyngeal samples positive for SARS CoV2. All the patients were confirmed local transmission cases from Karachi. Viral RNA extracted from both asymptomatic and symptomatic COVID-19 patients were included. Whole genome sequencing was performed as previously described through ONT MinIon platform. Briefly, cDNA was prepared from viral RNA using random primer mix (NEB, Massachusetts, United States), and LunaScript^®^ RT SuperMix Kit (Cat# E3010G). The cDNA was enriched using Q5 high-fidelity DNA polymerase (Cat #: M0492; NEB, USA) and ARTIC v3 primers (*Integrated DNA Technologies (ITD), Belgium*). The 400bp amplicon was used for library preparation after purification by AMPure XP beads (Agencourt Beckman Coulter™ A63881) and quantification with a Qubit fluorometerusing Qubit™ RNA BR Assay kit (Cat #, Q10210, ThermoFisher Scientific,Massachusetts, United States). For sequencing SQK-LSK104 ligation kit was used and samples were loaded as per standard protocol on a MinION MK1B instrument. All the samples were multiplexed in one run after barcoding. The samples were run for 72 hours on R9 flow cell. The acquired data was aligned in real time with SARS CoV2 reference sequence (NC_045512.2) using Min Know Files in SAM/BAM format were accessed by SAM tools v. 1.9-11.

### Variant calling and phylogenetic profiling

Variant calling was performed using BCF tools v. 1.9. Further variant annotation was done by ANNOVAR. The consensus sequences were generated by mapping the variants to the reference genomes using BCF tools followed by submission to the GISAID database and NCBI. The initial phylogenetic analysis was performed using 570 genome sequences retrieved from the GISAID, epicov™. The fast alignment was performed using MAFFT. Moreover, IQ-TREE v. 2.1.2 provided by Augur (github.com/nextstrain/augur) was used to infer maximum-likelihood trees. The generated tree was visualized using FigTree 1.4.3(http://tree.bio.ed.ac.uk/software/figtree). Viral clades were identified by nucleotide or amino acid substitutions and Nextstrain nomenclature (https://nextstrain.org/ncov) was followed for matching.

Genomic epidemiological analysis was performed using 564 complete sequences downloaded from GISAID database. Phylogenetic tree was constructed using NEXTSTRAIN (https://www.nextstain.org) via augur pipeline, after inclusion of the four sequences from the current study to the common FASTA file containing all downloaded sequences. The robustness of individual nodes was statistically determined by comparing with 1000 bootsteps replicates. The viral lineage was determined by using Phylogenetic Assignment of Named Global Outbreak LINeages tool (https://github.com/cov-lineages/pangolin).

### Statistical analysis

Median Ct values per month for Orf1ab gene were recorded from May to November 2020. For statistical analyses and graphs SPSS version 22 was used.

## Acknowledgement

The authors are thankful to Scientechnic for providing the MinION MK1B instrument and timely supply of all ordered consumable for this project during the pandemic.

## References

1. Zhou P, Yang X-L, Wang X-G, Hu B, Zhang L, et al. (2020) A pneumonia outbreak associated with a new coronavirus of probable bat origin. nature 579: 270–273.

2. Lu R, Zhao X, Li J, Niu P, Yang B, et al. (2020) Genomic characterisation and epidemiology of 2019 novel coronavirus: implications for virus origins and receptor binding. The Lancet 395: 565–574.

3. Xiao K, Zhai J, Feng Y, Zhou N, Zhang X, et al. (2020) Isolation of SARS-CoV-2-related coronavirus from Malayan pangolins. Nature: 1–4.

4. Huang C, Wang Y, Li X, Ren L, Zhao J, et al. (2020) Clinical features of patients infected with 2019 novel coronavirus in Wuhan, China. The lancet 395: 497–506.

5. Mercatelli D, Giorgi FM (2020) Geographic and Genomic Distribution of SARS-CoV-2 Mutations.

6. Wang C, Horby PW, Hayden FG, Gao GF (2020) A novel coronavirus outbreak of global health concern. The Lancet 395: 470–473.

7. Yoshimoto FK (2020) The Proteins of Severe Acute Respiratory Syndrome Coronavirus-2 (SARS CoV-2 or n-COV19), the Cause of COVID-19. The Protein Journal: 1.

8. Fung S-Y, Yuen K-S, Ye Z-W, Chan C-P, Jin D-Y (2020) A tug-of-war between severe acute respiratory syndrome coronavirus 2 and host antiviral defence: lessons from other pathogenic viruses. Emerging microbes & infections 9: 558–570.

9. Wu F, Zhao S, Yu B, Chen Y-M, Wang W, et al. (2020) A new coronavirus associated with human respiratory disease in China. Nature 579: 265–269.

10. Rahimi A, Mirzazadeh A, Tavakolpour S (2020) Genetics and genomics of SARS-CoV-2: A review of the literature with the special focus on genetic diversity and SARS-CoV-2 genome detection. Genomics.

11. van Dorp L, Acman M, Richard D, Shaw LP, Ford CE, et al. (2020) Emergence of genomic diversity and recurrent mutations in SARS-CoV-2. Infection, Genetics and Evolution: 104351.

12. Hodcroft EB, Hadfield J, Neher R, Bedford T (2020) Year-letter genetic clade naming for SARS-CoV-2 on Nextstain. org. Nextstrainorg June 2.

13. Hartley PD, Tillett RL, AuCoin DP, Sevinsky JR, Xu Y, et al. (2020) Genomic surveillance revealed prevalence of unique SARS-CoV-2 variants bearing mutation in the RdRp gene among Nevada patients. medRxiv.

14. Korber B, Fischer WM, Gnanakaran S, Yoon H, Theiler J, et al. (2020) Tracking changes in SARS-CoV-2 Spike: evidence that D614G increases infectivity of the COVID-19 virus. Cell 182: 812–827. e819.

15. Moore SC, Penrice-Randal R, Alruwaili M, Dong X, Pullan ST, et al. (2020) Amplicon based MinION sequencing of SARS-CoV-2 and metagenomic characterisation of nasopharyngeal swabs from patients with COVID-19. medRxiv.

16. GISAID (2020) Clade and lineage nomenclature aids in genomic epidemiology studies of active hCoV-19 viruses. Available from https://www.gisaid.org/references/statements-clarifications/clade-and-lineage-nomenclature-aids-in-genomicepidemiologyof-active-hcov-19-viruses/.

17. Yao H-P, Lu X, Chen Q, Xu K, Chen Y, et al. (2020) Patient-derived mutations impact pathogenicity of SARS-CoV-2. CELL-D-20-01124.

18. Pachetti M, Marini B, Benedetti F, Giudici F, Mauro E, et al. (2020) Emerging SARS-CoV-2 mutation hot spots include a novel RNA-dependent-RNA polymerase variant. Journal of Translational Medicine 18: 1–9.

19. Li Q, Wu J, Nie J, Zhang L, Hao H, et al. (2020) The impact of mutations in SARS-CoV-2 spike on viral infectivity and antigenicity. Cell 182: 1284–1294. e1289.

20. Bianchi M, Borsetti A, Ciccozzi M, Pascarella S (2020) SARS-Cov-2 ORF3a: Mutability and function. Int J Biol Macromol.

21. Lam J-Y, Yuen C-K, Ip JD, Wong W-M, To KK-W, et al. (2020) Loss of orf3b in the circulating SARS-CoV-2 strains. Emerging Microbes & Infections: 1–678.

22. Laamarti M, Alouane T, Kartti S, Chemao-Elfihri M, Hakmi M, et al. (2020) Large scale genomic analysis of 3067 SARS-CoV-2 genomes reveals a clonal geodistribution and a rich genetic variations of hotspots mutations. bioRxiv.

23. Hodcroft EB, Zuber M, Nadeau S, Comas I, Candelas FG, et al. (2020) Emergence and spread of a SARS-CoV-2 variant through Europe in the summer of 2020. MedRxiv.

24. Kamikubo Y, Hattori T, Takahashi A (2020) Epidemic trends of SARS-CoV-2 associated with immunity, race, and viral mutations.

25. Liu S, Shen J, Fang S, Li K, Liu J, et al. (2020) Genetic spectrum and distinct evolution patterns of SARS-CoV-2. Frontiers in Microbiology 11: 2390.

26. Bartolini B, Rueca M, Gruber CEM, Messina F, Giombini E, et al. (2020) The newly introduced SARS-CoV-2 variant A222V is rapidly spreading in Lazio region, Italy. medRxiv.

